# The *E. coli* molecular phenotype under different growth conditions

**DOI:** 10.1101/082032

**Authors:** Mehmet U. Caglar, John R. Houser, Craig S. Barnhart, Daniel R. Boutz, Sean M. Carroll, Aurko Dasgupta, Walter F. Lenoir, Bartram L. Smith, Viswanadham Sridhara, Dariya K. Sydykova, Drew Vander Wood, Christopher J. Marx, Edward M. Marcotte, Jeffrey E. Barrick, Claus O. Wilke

**Author notes:** Corresponding author (MUC); (EMM); (JEB); (COW).

## Abstract

Modern systems biology requires extensive, carefully curated measurements of cellular components in response to different environmental conditions. While high-throughput methods have made transcriptomics and proteomics datasets widely accessible and relatively economical to generate, systematic measurements of both mRNA and protein abundances under a wide range of different conditions are still relatively rare. Here we present a detailed, genome-wide transcriptomics and proteomics dataset of *E. coli* grown under 34 different conditions. Additionally, we provide measurements of doubling times and *in-vivo* metabolic fluxes through the central carbon metabolism. We manipulate concentrations of sodium and magnesium in the growth media, and we consider four different carbon sources glucose, gluconate, lactate, and glycerol. Moreover, samples are taken both in exponential and stationary phase, and we include two extensive time-courses, with multiple samples taken between 3 hours and 2 weeks. We find that exponential-phase samples systematically differ from stationary-phase samples, in particular at the level of mRNA. Regulatory responses to different carbon sources or salt stresses are more moderate, but we find numerous differentially expressed genes for growth on gluconate and under salt and magnesium stress. Our data set provides a rich resource for future computational modeling of *E. coli* gene regulation, transcription, and translation.

## Introduction

A goal of systems biology has been to understand how phenotype originates from genotype. The phenotype of a cell is determined by complex regulation of metabolism, gene expression, and cell signaling. Understanding the connection between phenotype and genotype is crucial to understanding disease and for engineering biology^1^. Computational models are particularly well suited to studying this problem, as they can synthesize and organize diverse and complex data in a predictive framework, but detailed experimental studies including many samples are needed to understand interactions between different types of omics data^2^. Much effort is currently being spent on understanding how to best integrate information collected about multiple cellular subsystems that is derived from different types of high-throughput measurements. For example, there are many proposed approaches for relating gene expression and protein abundances, focusing on integrative, whole-cell models^2–5^.

Given the growing interest in integrative modeling approaches, there is a pressing need for high quality genome-scale data that is comparable across cellular subsystems and reflects many different external conditions. *E. coli* is an ideal organism to study genome-wide, multi-level regulatory effects of external conditions, since it is well adapted to the laboratory environment^6^ and was one of the first organisms studied at the whole-genome level^7^. There have been a number of studies of the *E. coli* transcriptome and/or proteome in response to different growth conditions. For example, in cells growing at high density, expression of most amino acid biosynthesis genes is down-regulated and expression of chaperones is up-regulated, suggesting stresses that these cells experience^8^. Exposure of *E. coli* to reduced temperature leads to changes in gene-expression patterns consistent with reduced metabolism and growth^9^. Under long-term glucose starvation, mRNAs are generally down-regulated while the protein response is more varied^10^. Specifically, the copy numbers of proteins involved in energy-intensive processes decline whereas those of proteins involved in nutrient metabolism remain constant, likely to provide the cell with the ability to jump-start metabolism when nutrients become available again. A few other larger-scale studies have measured mRNA and/or protein abundances under multiple conditions^11–14^.

Here, we provide a systematic analysis of *E. coli* gene expression under a wide variety of different conditions. We measure both mRNA and protein abundances, at exponential and stationary phases, for growth conditions including different carbon sources and different salt stresses. We find that mRNAs and proteins display divergent responses to the different growth conditions. Further, growth phase yields more systematic differences in gene expression than does either carbon source or salt stress, though this effect is more pronounced in mRNAs than in proteins. We expect that our data set will provide a rich resource for future modeling work.

## Results

### Experimental design and data collection

We grew multiple cultures of *E. coli* REL606, from the same stock, under a variety of different growth conditions. We measured RNA abundances under all conditions and matching protein abundances for approximately 2/3 of the conditions (Figure 1 and Supplementary Table S1). We also measured central metabolic fluxes for a subset of conditions using glucose as carbon source. Results from one of these conditions, long-term glucose starvation, have been presented previously^10^. Conditions not previously described include one additional starvation experiment, using glycerol instead of glucose as carbon source, exponential and stationary phase cultures using either gluconate or lactate as carbon source, and conditions varying Mg^2+^ and Na^+^ concentrations.

**Figure 1:**
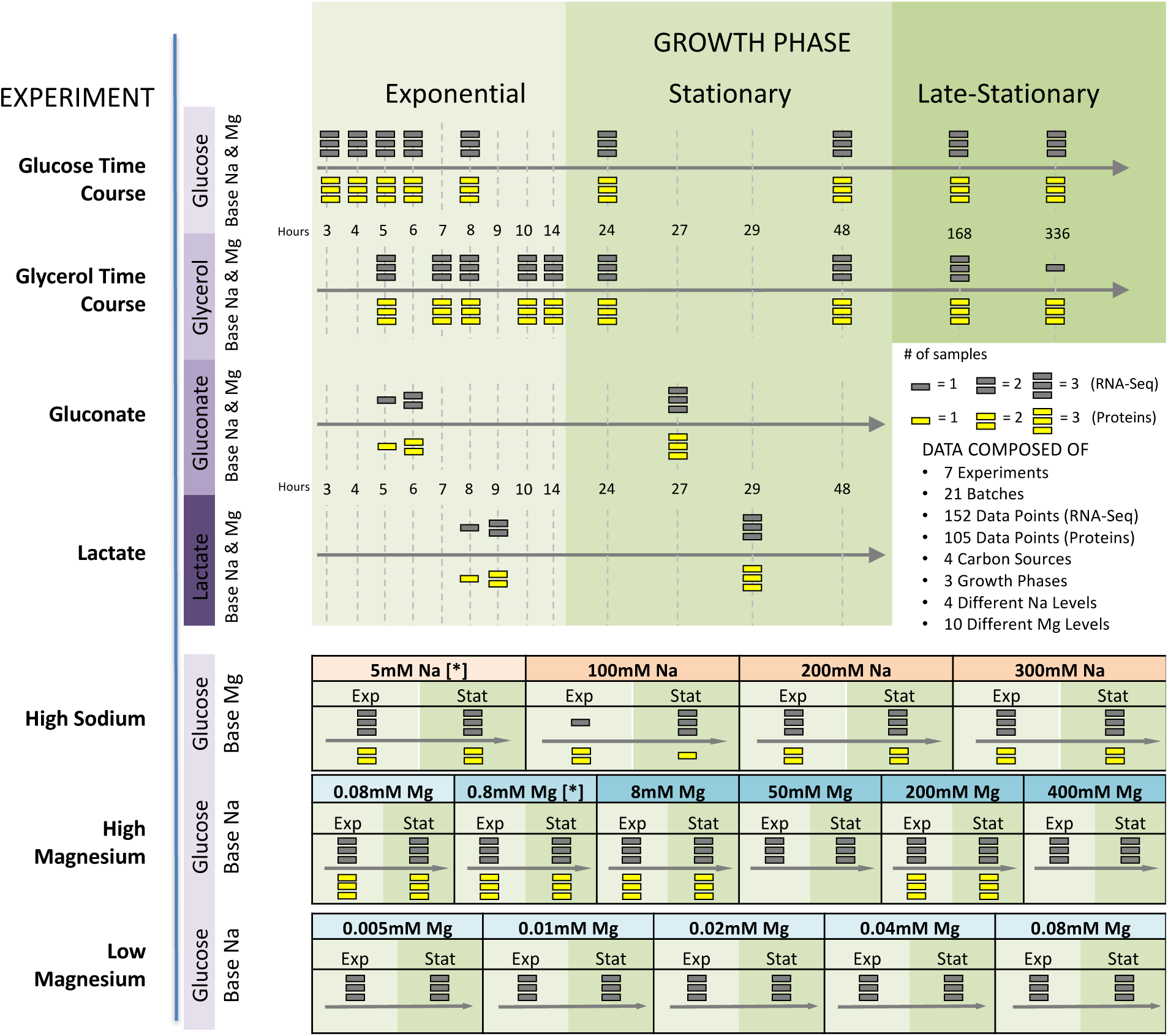
Experimental setup. We performed seven different experiments in which we varied the duration of growth and the temporal density of sampling, the carbon source, and ion concentrations. For each experimental condition, bacteria were grown in three biological replicates. Conditions that are listed multiple times represent independent replicates of those conditions. We subsequently performed whole-transcriptome RNA-Seq for all experimental conditions and mass-spec proteomics for the majority of them. (No proteomics was performed for the low-magnesium experiment.) We considered four different carbon sources: glucose, glycerol, gluconate, and lactate; we also considered high sodium and both low and high magnesium levels. For the time-course and carbon-source experiments, we used base-level Na^+^ (5 mM) and Mg^2+^ (0.8 mM) throughout (indicated by [*] in the sodium and magnesium experiments).

Measurements of RNA and protein abundances were carried out as previously described^10^. All resulting data sets were checked for quality, normalized, and log-transformed. Our final data set consisted of 152 RNA samples, 105 protein samples, and 65 flux samples (Supplementary Table S1). 59 of the flux samples are associated with high Mg^2+^ and high Na^+^ experiments.

Our raw RNA-seq and protein data covers 4196 distinct mRNAs and proteins, and our flux data covers 13 different metabolic reactions. All raw data files are available in appropriate repositories (see Methods for details), and final processed data are available as Supplementary Tables S2, S3, and S4.

Finally, we measured doubling times in exponential phase for all experimental conditions (Supplementary Table S5). We found that doubling times varied between 50 and 100 minutes among the various conditions (Figure 2). Growth was the fastest when glucose was used as carbon source and the slowest when the carbon source was lactose. Growth was also reduced for high Na^+^ concentrations and very high or low Mg^2+^ concentrations. Surprisingly, we found a broad range of Mg^2+^ concentrations (0.02mM to 200mM) in which growth rate remained virtually unchanged (Figure 2).

**Figure 2:**
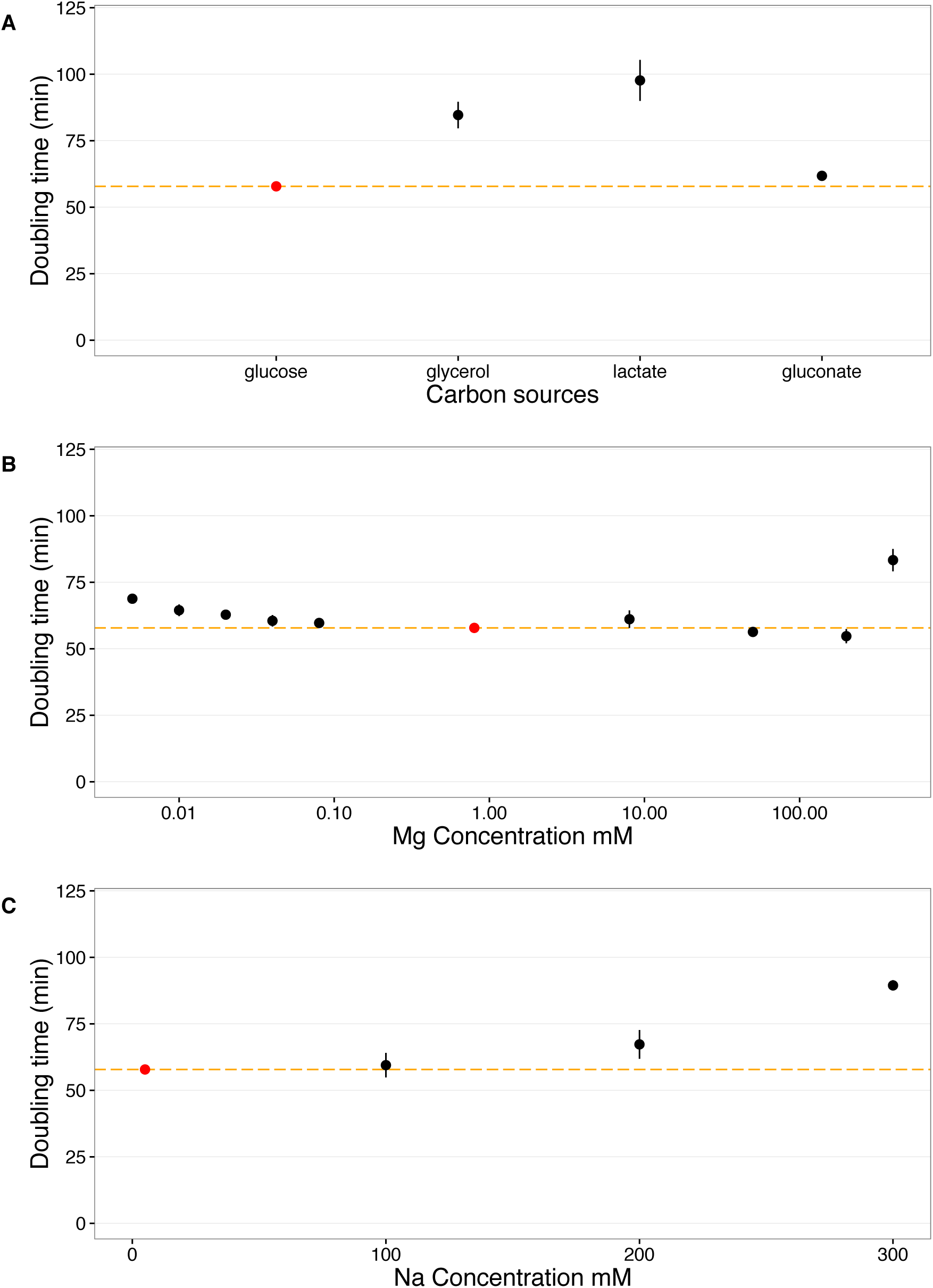
Doubling times under various growth conditions. We measured doubling times under exponential phase for all growth conditions. The red points and dashed orange lines represent the doubling time at the base condition (glucose, 5 mM Na^+^, 0.8 mM Mg^2+^). Doubling times were measured in triplicates and error bars represents 95% confidence intervals of the mean. (A) Doubling times with respect to carbon sources. (B) Doubling times with respect to Mg^2+^ concentrations. (C) Doubling times with respect to Na^+^ concentrations.

### Broad trends of gene expression differ between mRNA and proteins

To identify broad trends of gene expression among the different growth conditions, we performed hierarchical clustering on both mRNA and protein abundances (Figures 3 and 4). For mRNA, we found that differences in gene expression were primarily driven by the growth phase (exponential vs. stationary/late stationary). Nearly all exponential samples clustered together in one group, separate from the vast majority of stationary and late-stationary samples (Figure 3). Mg^2+^ levels, Na^+^ levels, and carbon source had less influence on the clustering results. Results were different for protein abundances (Figure 4), where growth phase had little effect on the clustering and instead samples seemed to group together by Na^+^ levels and carbon source.

**Figure 3:**
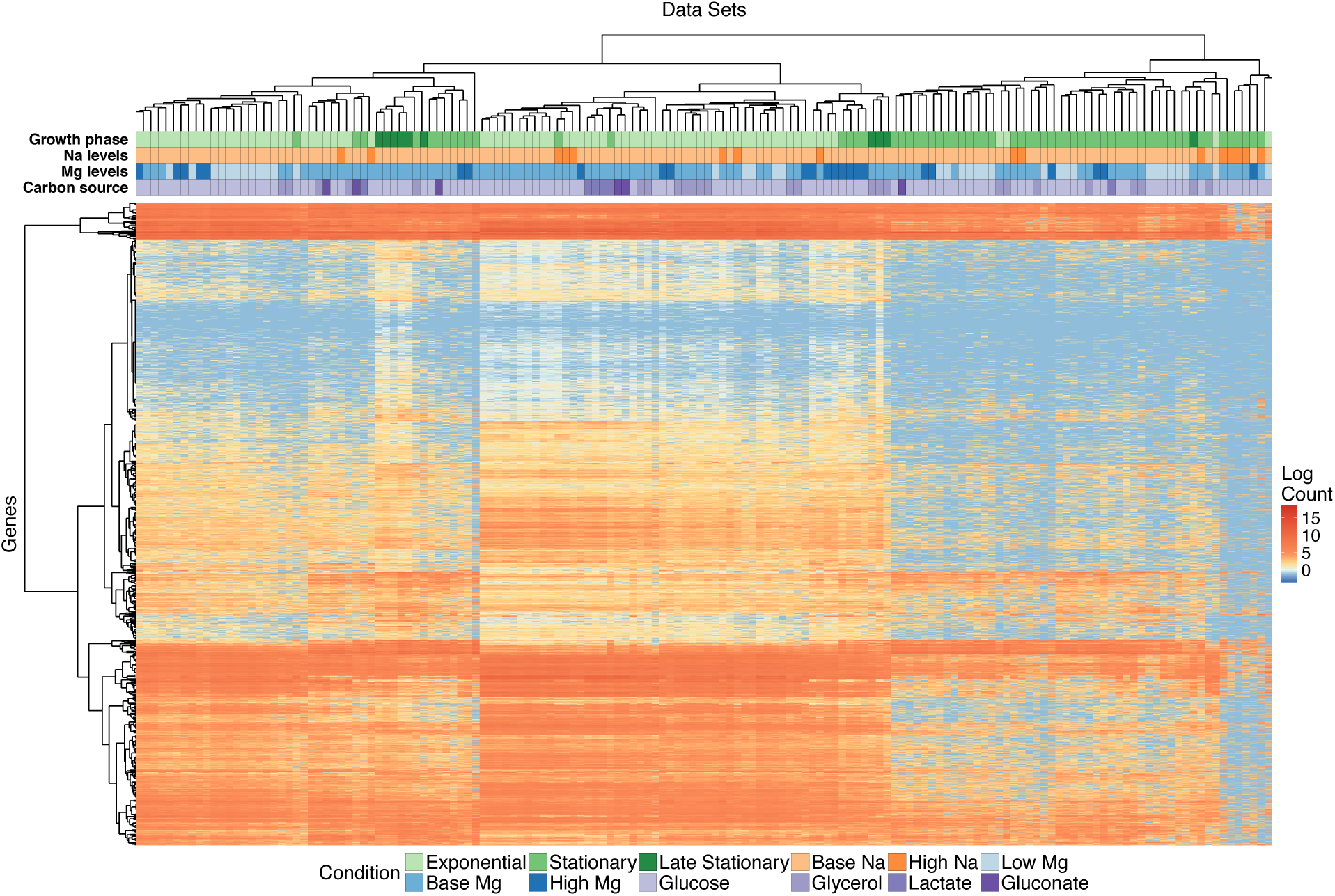
Clustering of mRNA abundances. The heatmap shows 4196 mRNA abundances for each of 152 samples, clustered both by similarity across genes and by similarity across samples. The growth conditions for each sample are indicated by the color coding along the top of the heatmap; the color coding is defined in the legend at the bottom.

**Figure 4:**
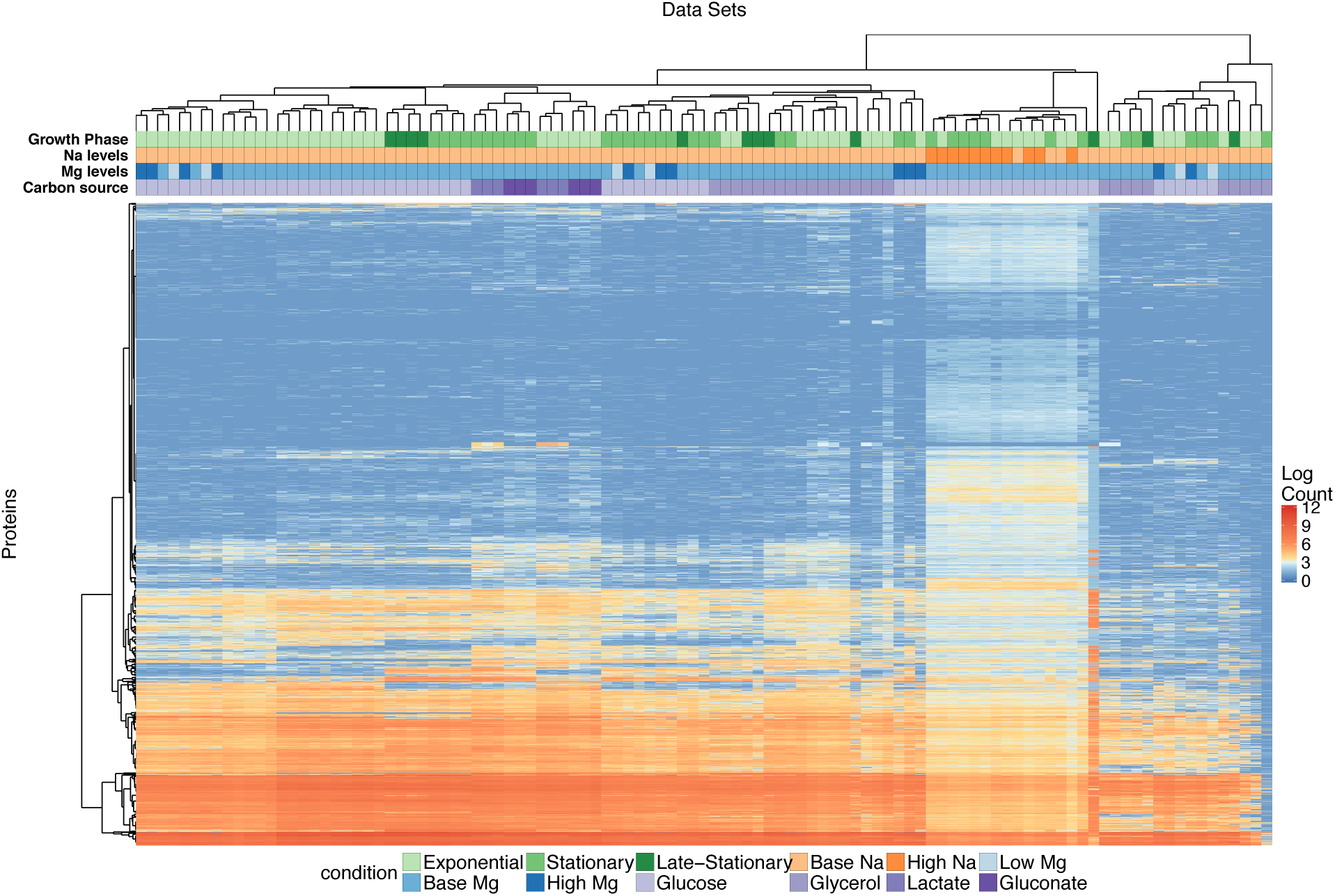
Clustering of protein abundances. The heatmap shows 4196 protein abundances for each of 105 samples, clustered both by similarity across genes and by similarity across samples. The growth conditions for each sample are indicated by the color coding along the top of the heatmap; the color coding is defined in the legend at the bottom.

To quantify the clustering patterns of mRNA and protein abundances, we defined a metric that measured how strongly clustered a given variable of the growth environment (growth phase, Mg^2+^ level, Na^+^ level, carbon source) was relative to the random expectation of no clustering. For each variable, we calculated the mean cophenetic distance between all pairs corresponding to the same condition (e.g., for growth phase, all pairs sampled at exponential phase and all pairs sampled at stationary/late stationary phase). The cophenetic distance is defined as the height of the dendrogram produced by the hierarchical clustering from the two selected leafs to the point where the two branches merge, and it is widely used to quantify how closely related any two leafs are in a clustering dendogram^15,16^. We generated null distributions of cophenetic distances under the assumption of no clustering by resampling mean cophenetic distances from dendograms with reshuffled leaf assignments, and we then converted each observed mean cophenetic distance into a z-score using the mean and variance of the corresponding null distribution. Thus, we carried out a non-parametric, random permutation test where we determined the null-distribution of our test statistic by resampling. A z-score below −1.96 indicates that the mRNA or protein abundances are clustered significantly by the corresponding variable.

We found that mRNA abundances were significantly clustered by growth phase, with a z-score of −30.98, and by Mg^2+^ level, with a z-score of −3.21 (Table 1). The z-scores for Na^+^ level and carbon source were −1.89 and 1.21, respectively, which are not significantly different from zero. Moreover, when we calculated a z-score for batch number, we found that batch effects did not significantly influenced mRNA abundances, with *z* = −1.43. Batch numbers represent cultures grown at the same time, in parallel.

**Table 1:** Clustering of mRNA and protein abundances by different growth conditions. The *z* scores represent mean cophenetic distances between all pairs of conditions with the same label, normalized by the distribution of mean distances obtained after randomly reshuffling condition labels. The overall *z* score tests for significant clustering within a given variable, and the individual *z* score tests for significant clustering within a given condition. Significant clustering (defined as |*z*| >2) is indicated with a *. The *z*-scores of individual batches are provided in Supplementary Tables S6 and S7.

| mRNA |  |  |  |  |
| --- | --- | --- | --- | --- |
| Variable | Overall z score | Condition | z score | # samples |
| Growth phase | -30.98 * | Exponential | -15.49 * | 79 |
|  |  | Stationary | -0.94 | 63 |
|  |  | Late stationary | -2.9 * | 10 |
| Carbon source | 1.21 | Glucose | 1.19 | 115 |
|  |  | Glycerol | -0.48 | 25 |
|  |  | Lactate | -1.93 | 6 |
|  |  | Gluconate | -0.46 | 6 |
| Mg Level | -3.21 * | Low Mg<sup>2+</sup> | 0.38 | 36 |
|  |  | Base Mg<sup>2+</sup> | -2.21 * | 92 |
|  |  | High Mg<sup>2+</sup> | -0.97 | 24 |
| Na Level | -1.89 | Base Na<sup>+</sup> | -1.83 | 136 |
|  |  | High Na<sup>+</sup> | 1.15 | 16 |
| Batch number | -1.43 |  |  |  |

| Protein |  |  |  |  |
| --- | --- | --- | --- | --- |
| Variable | Overall z score | Condition | z score | # samples |
| Growth phase | -1.27 | Exponential | -0.82 | 56 |
|  |  | Stationary | 0.23 | 37 |
|  |  | Late stationary | -0.08 | 12 |
| Carbon source | -2.79 * | Glucose | -2.33 * | 66 |
|  |  | Glycerol | 1.35 | 27 |
|  |  | Lactate | -2.73 * | 6 |
|  |  | Gluconate | -2.63 * | 6 |
| Mg Level | -0.50 | Low Mg<sup>2+</sup> | 0.85 | 6 |
|  |  | Base Mg<sup>2+</sup> | -0.44 | 87 |
|  |  | High Mg<sup>2+</sup> | -0.42 | 12 |
| Na Level | -1.74 | Base Na<sup>+</sup> | -0.94 | 94 |
|  |  | High Na<sup>+</sup> | -5.61 * | 11 |
| Batch number | -20.54 * |  |  |  |

For protein abundances, the variable carbon source was significantly clustered, with a z-score of −2.79, and the other variables Na^+^ levels, growth phase, and Mg^2+^ levels were not significantly clustered, with z scores of −1.74, −1.27, and −0.5, respectively (Table 1). Batch number had a z-score of −20.54, which implies that there were strong batch effects present in the protein data. In general, batch effects may represent fluctuations in incubator temperatures, slight differences in growth medium composition or water quality, effects of reviving the initial inoculum of cells, or effects of sample preparation and analysis, among other possibilities. Here, since batch effects were so pronounced in the proteomics data and not in the mRNA data, we suspect that they were primarily caused by proteomics sample preparation and analysis.

In summary, mRNA abundances were clustered by growth phase and Mg^2+^ levels, whereas protein abundances were clustered by carbon source. Protein abundances were also strongly influenced by batch effects, unlike the mRNA data (Table 1, Supplementary Table S6, S7).

### Identification of differentially expressed genes

We next asked under which conditions and to what extent RNA and protein expression were altered. To identify differentially expressed mRNAs and proteins, we used DESeq2^17^. Since a detailed comparison of exponential vs. stationary phase has been published previously for the glucose time-course experiment^10^, here we focused on differences among ion concentrations or carbon sources within either exponential or stationary phase.

For each growth phase, we defined the base level reference condition to be growth in glucose with 5 mM Na^+^ and 0.8 mM Mg^2+^. This is the baseline formulation of media used in the glucose time-course samples^10^. We then compared RNA and protein abundances between this reference condition and the alternative conditions (different carbon sources, elevated Na^+^, and elevated or reduced Mg^2+^) separately for each growth phase (Supplementary Table S8).

We defined significantly differentially expressed genes as those whose abundance had at least a two-fold change (log_2_ fold change > 1) between the reference condition and a chosen experimental condition, at a false-discovery-rate (FDR) corrected *P* value < 0.05. We found that the number of significantly differentially expressed mRNAs and proteins varied substantially between exponential and stationary phase and between mRNAs and proteins (Figure 5 and (Supplementary Table S9). In general, there were fewer differentially expressed genes in stationary phase than in exponential phase. Further, protein abundances showed the most differential regulation for high Na^+^ and for the carbon sources glycerol and lactate, whereas mRNA showed the most differential regulation for high Na^+^ levels in stationary phase, and for low Mg^2+^ levels and for the carbon sources glycerol and lactate in exponential phase (Figure 5).

**Figure 5:**
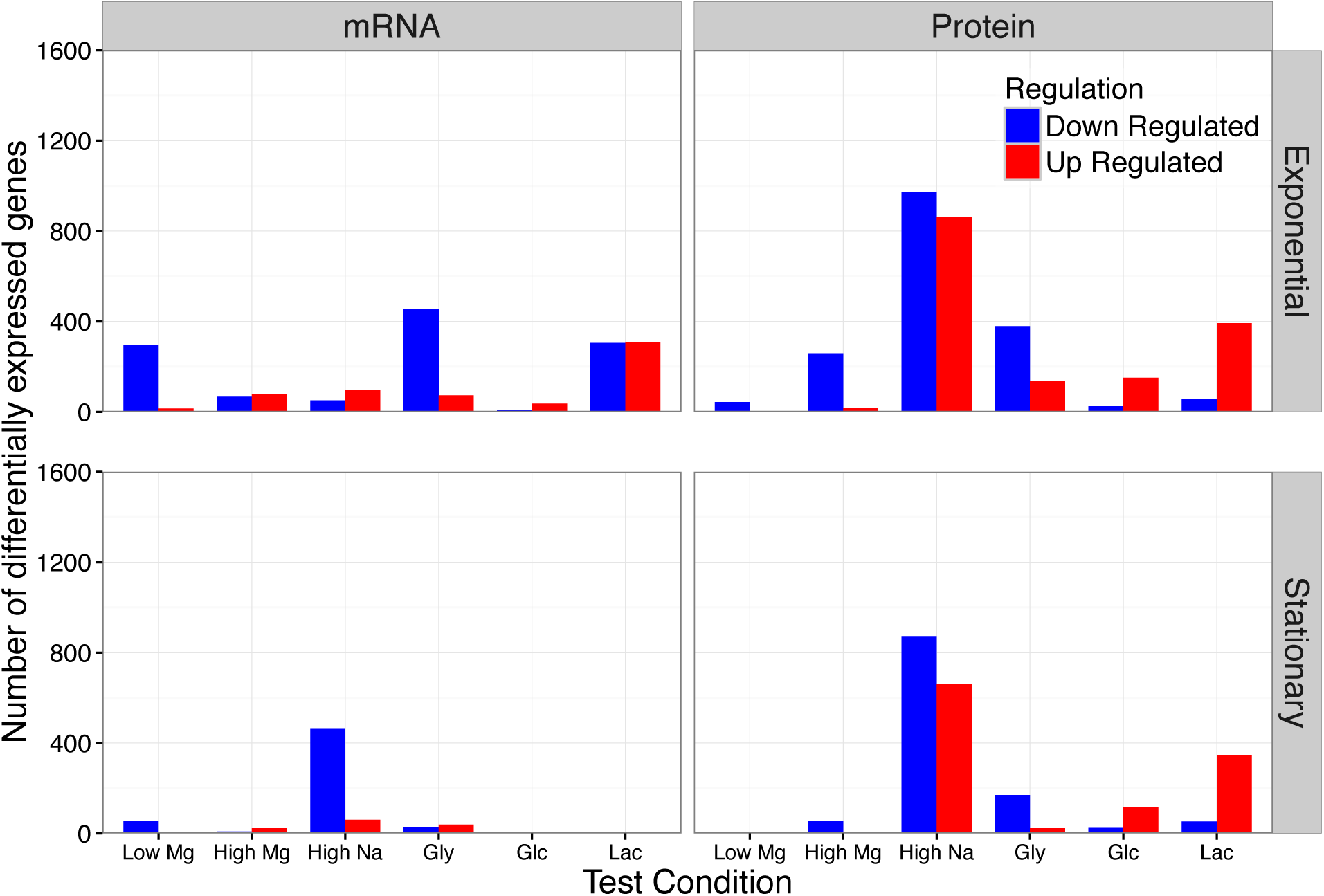
Number of differentially expressed genes under different conditions. We separately analyzed mRNA and protein abundances, each for both exponential and stationary growth phase. In all four cases, gene expression levels were compared to the corresponding condition with glucose as carbon source and baseline sodium and magnesium levels. Differentially expressed genes were defined has having at least a two-fold change relative to baseline and a false-discovery rate <0.05. Carbon sources are abbreviated as follows: “Gly”: glycerol, “Glc”: gluconate, “Lac”: lactate.

Next, we asked how much overlap there was among differentially expressed genes between the various growth conditions. To simplify this analysis, we did not distinguish between up- or down-regulated genes, and we combined low and high Mg^2+^ into one group “Mg stress” and glycerol, lactate, and gluconate into one group “carbon source”. (Note that differentially expressed genes were still identified for individual conditions, as described above, and were combined into “Mg stress” and “carbon source” only for the final comparison.) At the mRNA level, there was some overlap (21.7%) between carbon source and Mg^2+^ stress in exponential phase. All other overlaps where minimal, ~5% or less (Figure 6). At the protein level, there was overlap between Na^+^ stress and carbon source (15.6% in exponential phase, 10.7% in stationary phase), while all other overlaps were also minimal, ~5% or less (Figure 6).

**Figure 6:**
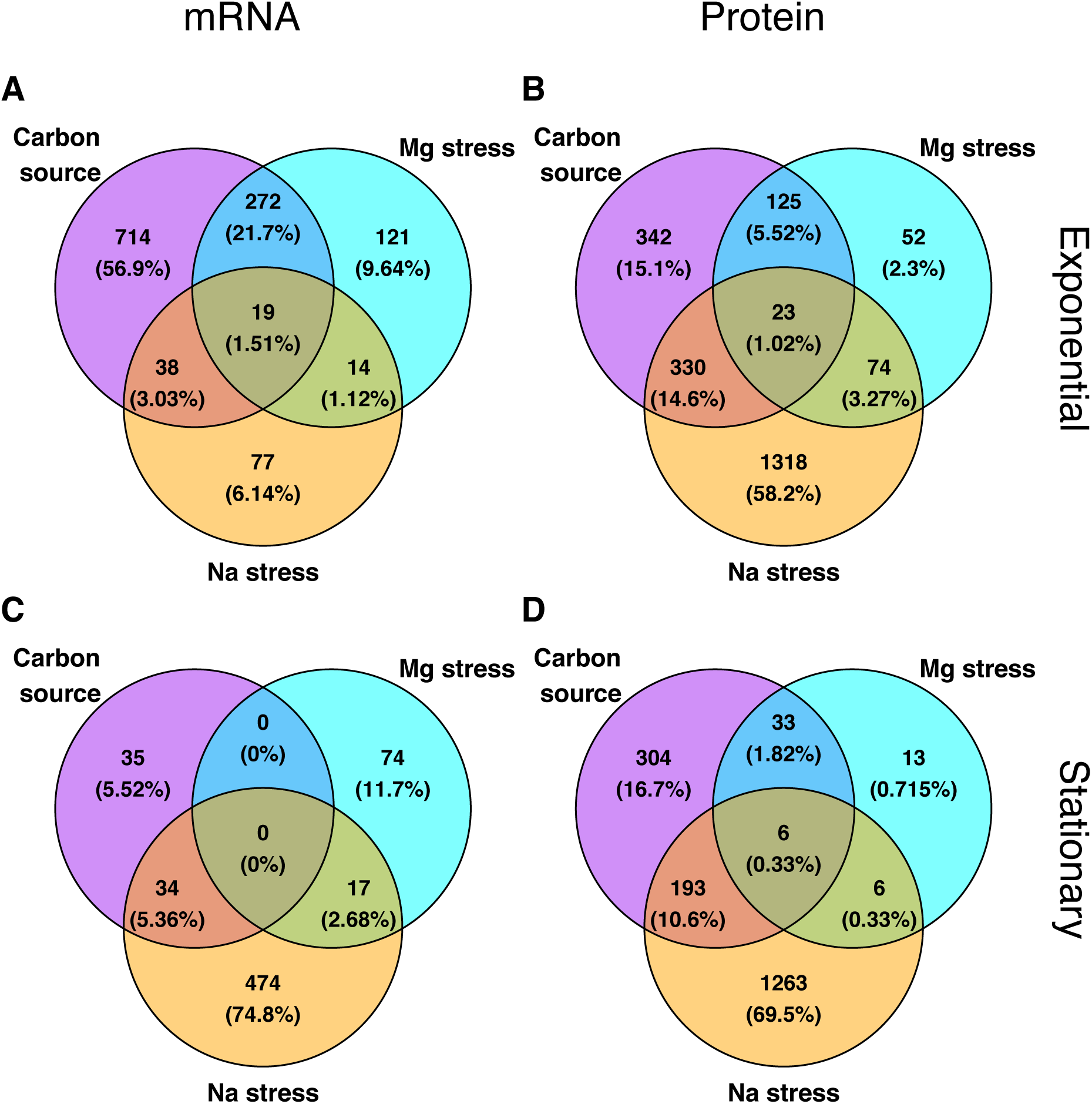
Overlap of differentially expressed genes among conditions. For all differentially expressed genes (identified as in Figure 5), we determined to what extent they were unique to specific conditions or appeared in multiple conditions. For simplicity, we here lumped all carbon-source experiments, all sodium experiments, and all magnesium experiments into one group each. Overall, we found relatively little overlap in the differentially expressed genes among these conditions. (A) mRNA, exponential phase. (B) protein, exponential phase. (C) mRNA, stationary phase. (D) protein, stationary phase.

We also identified significantly altered biological pathways and molecular activities of gene products (Supplementary Table S10). We used the Kyoto Encyclopedia of Genes and Genomes (KEGG)^18^ for biological pathways and annotations from the Gene Ontology (GO) Consortium for molecular functions^19^. Figure 7 and Supplementary Figure 1 show the top 5 significantly altered biological pathways (as defined in the KEGG database) and molecular functions (as defined by GO annotations) under different conditions, respectively, as determined by DAVID^20^. In all cases, we used a cutoff of 0.05 on false-discovery-rate (FDR)-corrected *P* values to identify significant annotations. We found numerous significantly altered KEGG pathways (Figure 7) molecular functions (Supplementary Figure S1).

**Figure 7:**
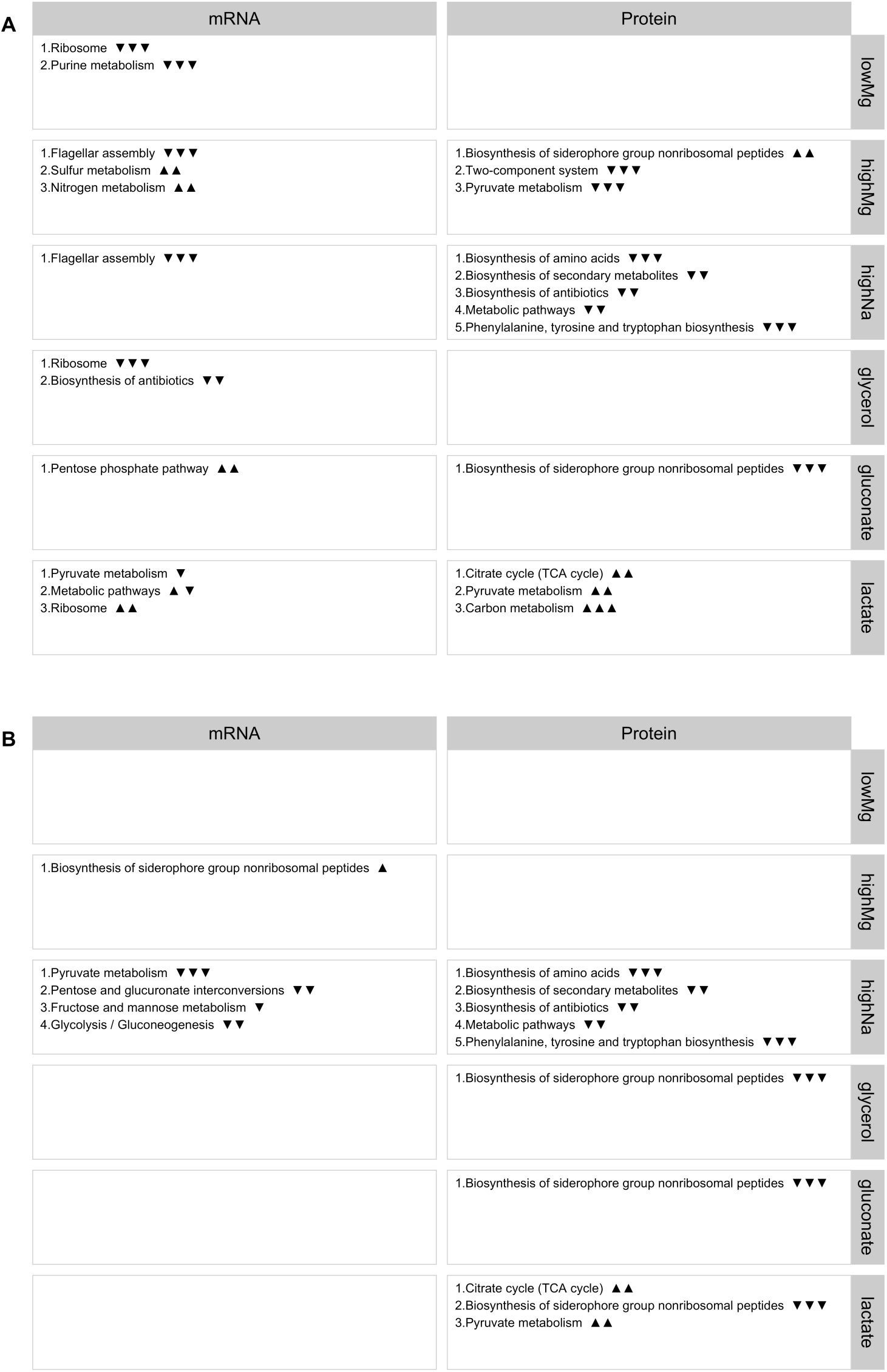
Significantly differentially expressed KEGG pathways. For each condition, we show the top-5 differentially expressed KEGG pathways as determined by either mRNA or protein abundances. Empty boxes indicate that no differentially expressed pathways were found. The arrows next to pathway names indicate the proportion of up- and down-regulated genes among the significantly differentially expressed genes in this pathway. One up arrow indicates that 60% or more of the genes are up-regulated, two arrows correspond to 80% or more genes, and three arrows correspond to 95% or more genes being up-regulated. Similarly, down arrows indicate the proportion of down-regulated genes. (A) Exponential phase. (B) Stationary phase.

In addition to identifying altered pathways and molecular activities, we identified the individual, most highly differentially expressed genes associated with specific pathways and/or functions (Supplementary Figures S2-S33). As an example, the differentially expressed mRNAs associated with significantly altered KEGG pathways under high Mg^2+^ concentrations in exponential phase are shown in Figure 8A. Three pathways are significantly altered; sulfur metabolism and nitrogen metabolism are mostly up-regulated and flagellar assembly is mostly down-regulated. Changes in sulfur metabolism in this condition might reflect a linked increased in the concentration of sulfate (SO_4_^2−^), as this was the counterion in the salt that was added to increase Mg^2+^ levels. By contrast, using lactate instead of glucose as carbon source caused up-regulation of pyruvate metabolism, citrate cycle, and carbon metabolism at the protein level in exponential phase (Figure 8B).

**Figure 8:**
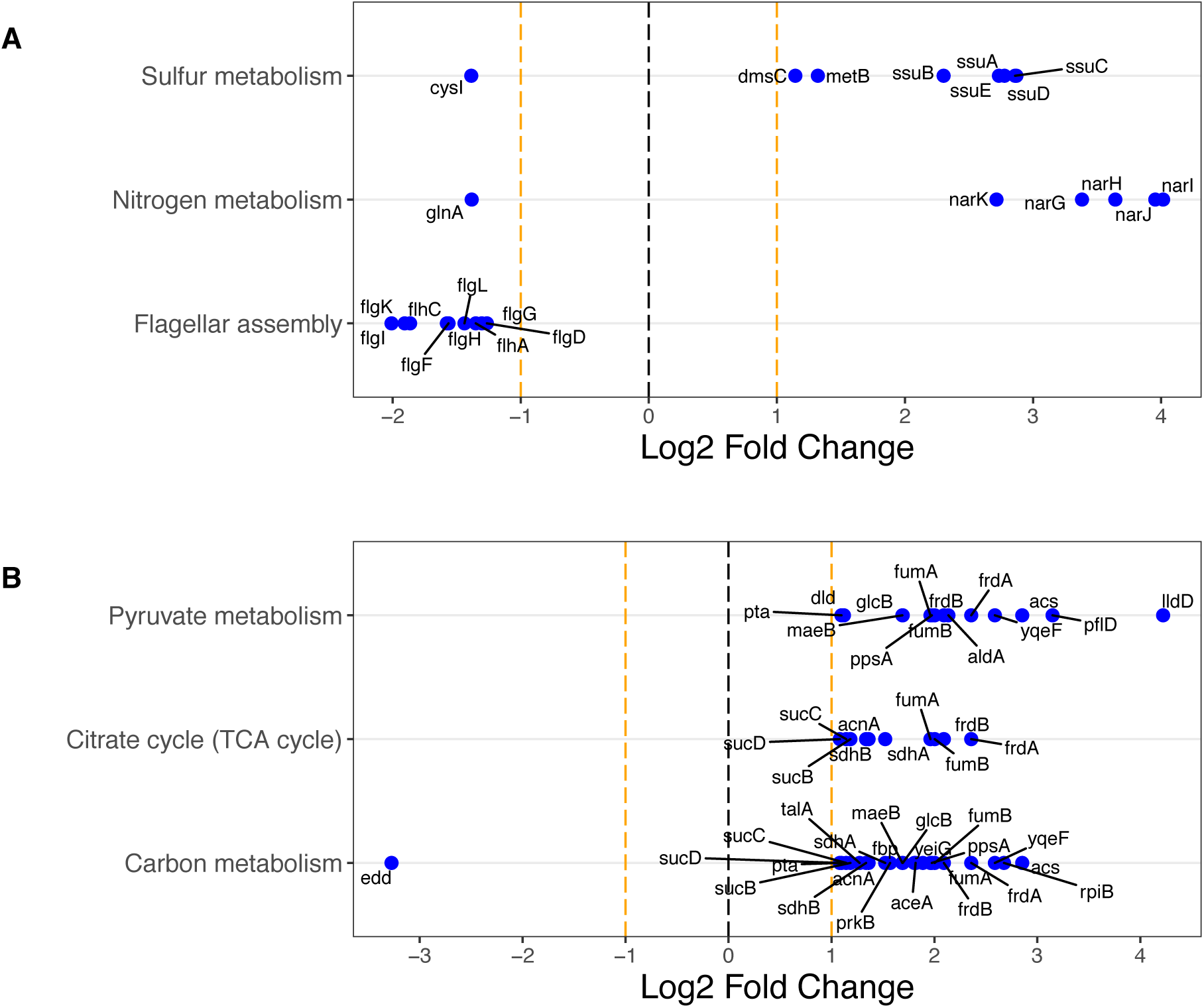
Examples of significantly differentially expressed KEGG pathways and associated genes. The top differentially expressed KEGG pathways are shown along the *y* axis, and the relative fold change of the corresponding genes is shown along the *x* axis. For each KEGG pathway, we show up to 10 of the most significantly changing genes. (A) Differentially expressed mRNAs under high Mg^2+^ levels in exponential phase against the control of samples at base Mg^2+^ level in exponential phase. (B) Differentially expressed proteins under lactate as carbon source in exponential phase against the control of samples with glucose as carbon source in exponential phase. Significant changes for all conditions are shown in Supplementary Figures 2-33.

Finally, we asked to what extent differentially expressed genes might be determined by bacterial growth rate, as measured by doubling time. We repeated our DeSeq2 analyses but included in our design formula a term representing the doubling time (see Methods). We found that in general, differences in these analyses are small; the most significantly changed genes when not controlling for doubling time are the most significantly changed genes when controlling for doubling time (Supplementary Table S9). One major exception were protein abundances in response to different carbon sources. In this scenario, many new genes appeared when controlling for doubling time, both in terms of the relative proportion of genes found and in terms of absolute numbers (Supplementary Figure S34, Supplementary Table S11). We identified the significantly altered pathways associated specifically with those genes, and we found that the top hits were related to biosynthesis in both exponential and stationary growth phases (Supplementary Table 12).

### Metabolic flux ratios under salt stress

For the high sodium and high magnesium experiments, we also determined metabolic flux through central metabolism by analyzing^13^ C incorporation into protein-bound amino acids. We here analyzed only flux samples taken in exponential phase, since stationary-phase samples have an unclear interpretation^10^. For each condition, flux samples were analyzed in triplicate (except one, which was analyzed in duplicate only), and 13 different flux ratios were measured for each sample. The flux ratios were then averaged across replicates (Supplementary Figure S35). We saw no significant changes in flux ratios with increasing Na^+^ (linear regression, all *P* > 0.05 after FDR correction, Supplementary Table S13). Results were similar for Mg^2+^. Due to the wide range of Mg^2+^ concentrations considered, we regressed flux ratios against log-transformed Mg^2+^ concentrations. Again, we saw no significant changes in any flux ratio with increasing Mg^2+^ (linear regression, all *P* > 0.05 after FDR correction, Supplementary Table S13).

We also asked whether the flux ratios changed with doubling time rather than with ion concentration, since doubling time is not necessarily monotonic in ion concentration (Figure 2B). For this analysis, we pooled all flux measurements and plotted flux ratios against doubling times (Figure 9). Again, we saw no significant relationship between flux ratios and doubling time after FDR correction (Supplementary Table S14). However, we note that the branches erythrose-4-phosphate from pentose-5-phosphate and pyruvate from malate (upper bound) showed a significant relationship before correction for multiple testing (*P* = 0.026 and *P* = 0.018, respectively, Supplementary Table S14), both driven by one outlying data point for the slowest-growing condition, at 300 mM Na^+^.

**Figure 9:**
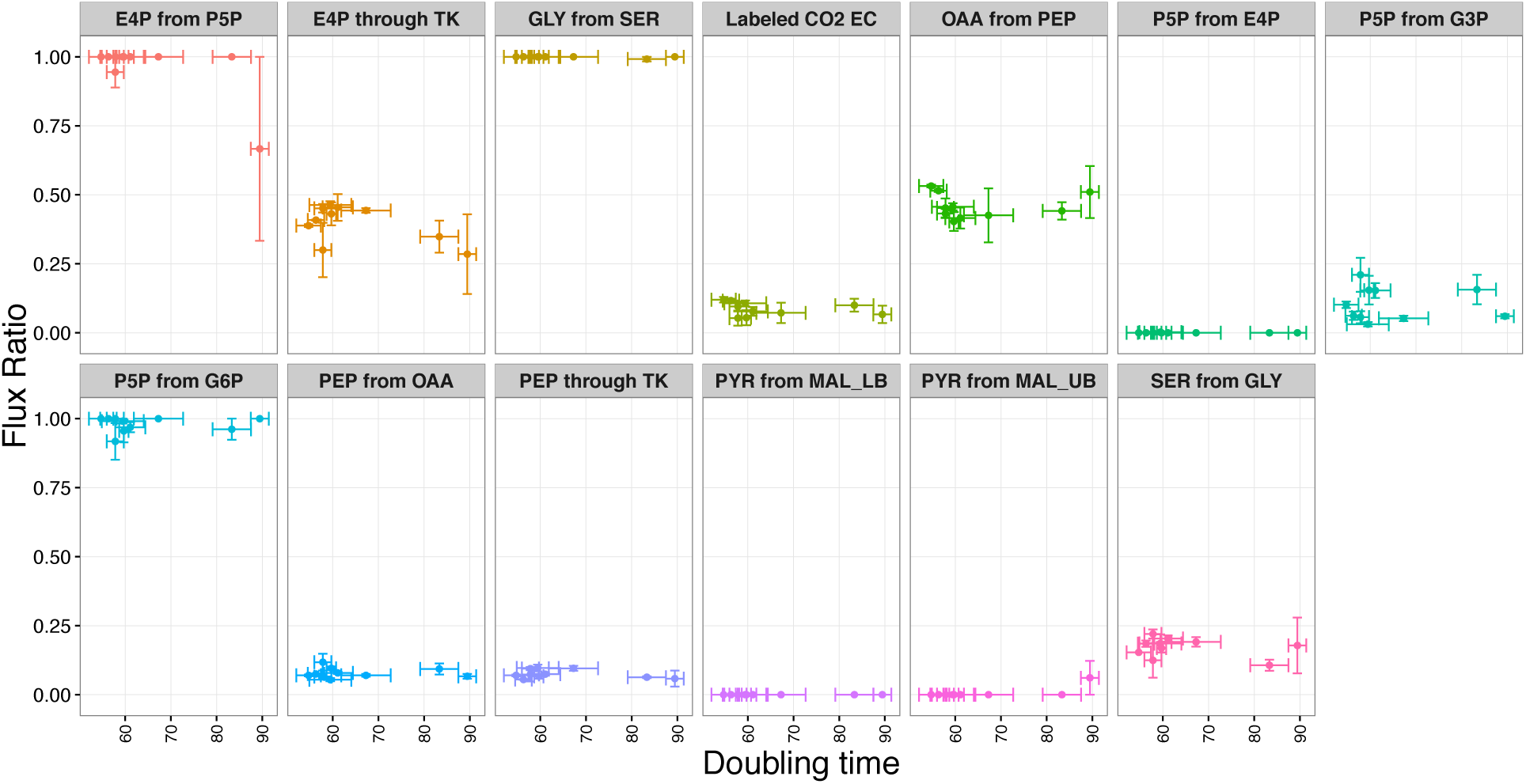
Flux ratios versus doubling times. 13 different flux ratios were measured for varying Na^+^ and Mg^2+^ concentrations (Supplementary Figure 34). Here, these flux ratios are shown as a function of the corresponding doubling times. The specific fluxes considered and their shorthand labels as used here are defined in Ref. 35. There was no significant association between any of the flux ratios and doubling time after correction for multiple testing (Supplementary Table 14).

## Discussion

We studied the regulatory response of *E. coli* under a wide variety of different growth conditions. The experimental conditions we considered include four different carbon sources, different levels of Na^+^ and Mg^2+^ stress, and growth into deep stationary phase, up to two weeks post inoculation. We found that gene regulation changes the most with respect to growth phase; in general, the exponential phase under one condition is more similar to the exponential phase under another condition than to the stationary phase under the same condition. Further, we found little overlap in differentially expressed genes under different growth conditions. Finally, we found that the ratios of fluxes through alternative branches within central metabolism remained approximately constant under salt stress, despite substantial changes in doubling times.

Our data provides a comprehensive picture of *E. coli* in terms of number, range, and depth of different stresses, comparable and complementary to other recently published datasets. For example, Schmidt *et al.*^12^ considered 22 unique conditions and measured abundances of >2300 proteins. mRNA abundances were not measured. Soufi *et al*.^11^ considered 10 unique conditions and also measured abundances of >2300 proteins. They were interested primarily in up- and down-regulated proteins under different ethanol stresses, and they found down-regulation of genes associated with ribosomes and protein biosynthesis during ethanol stress. Such genes were similarly down-regulated in our study during stress induced by high Na^+^ concentrations. Lewis *et al*.^13^ considered only 3 different carbon sources but measured mRNA and protein abundances in different strains adapted to these growth conditions. Finally, Lewis *et al*.^14^ compiled a database of 213 mRNA expression profiles covering 70 unique conditions, including different carbon sources, terminal electron acceptor, growth phase, and genotype. In comparison, we considered 34 unique conditions, measured 152 mRNA expression profiles, 105 protein expression profiles, and 59 flux profiles, and used the exact same *E. coli* genotype throughout.

Similar to our prior study^10^, we observed clear trends in the differential expression of mRNAs and proteins. In particular, we had reported previously^10^ that mRNAs are widely down-regulated in stationary phase whereas only select proteins are down-regulated. Consistent with that observation, we found here that mRNAs were significantly and strongly clustered by growth phase (*z* = −30.98) whereas proteins were not (*z* = −1.27). By contrast, at the protein level we saw significant clustering by carbon source (*z* = −2.79), which we did not see at the mRNA level. More specifically, we had found earlier^10^ that energy-intensive processes were down-regulated and stress-response proteins up-regulated in stationary phase. Similarly, we observed here that high Na^+^ stress conditions also led to the down-regulation of energy-intensive processes.

A number of genes and pathways that we found to be influenced by treatment conditions are consistent with prior knowledge from the literature. For instance, we found that increasing the concentration of Na^+^ and Mg^2+^ decreased transcription of the flagellar genes during exponential growth, as seen previously^21^. We also found that high concentrations of Mg^2+^ induce an increase in mRNA expression of sulfur and nitrogen transport proteins, and an increase in the enzymes necessary to produce the siderophore enterobactin (necessary for obtaining iron from the environment). These regulatory changes could be due to the high Mg^2+^ concentrations interfering with the bacterial membrane potential, and thereby inhibiting cotransporters that are coupled to this gradient. This effect has been previously described for iron^22^. High Na^+^ concentrations also significantly reduced the expression of a large number of proteins, mostly either involved in the biosynthesis of amino acids or components of the ribosome. These changes may simply reflect stress induced by the high Na^+^ concentrations used in these experiments.

Altering the carbon source, as well, provided predictable changes in gene expression. For instance, providing glycerol as the sole carbon source instead of glucose increases expression of *glpX,* part of the *glp* operon, which is involved in glycerol uptake^23^. Gluconate as a carbon source increases expression of genes from the *gnt* and *idn* operons, both involved in gluconate metabolism^24,25^. Finally, using lactate as a carbon source induces the expression of *IIdD (IctD),* a gene required for lactate utilization in *E. coli^26^.*

Large-scale, high-throughput gene-expression studies are frequently confounded by batch effects that can give rise to incorrect conclusions if they are not accounted for^27^. We saw such effects in our study as well. In our data, the batch number indicates bacterial samples that were grown at the same time. Not unexpectedly, our data showed significant clustering by batch number in the protein data, though not in the mRNA data (z scores of −20.54 and −1.43, respectively). Batch effects are not inherently a problem, as long as one is aware of their existence and analyzes data accordingly. Here, in our differential expression analysis, we corrected for batch effects by including batch as a distinct variable in the DESeq model (see Methods), as recommended by the DeSeq2 manual^28^. How to best correct for batch effects is a topic of ongoing investigations, and increasingly sophisticated methods are being developed to separate batch effects from real signal^29–32^. In particular, a recently developed semi-supervised normalization pipeline could be used to further investigate batch effects in this and other datasets^33^.

Given the many cellular changes observed in mRNA and protein levels, we turned to^13^ C labeling techniques^10,34,35^ to examine the extent to which these changes affected the relative flux of metabolites through different central metabolic pathway branch points during exponential growth. For this work, we concentrated upon growth on glucose during Na^+^ and Mg^2+^ stresses. Across these conditions, growth rates change over nearly a two-fold range, with the doubling time changing from approximately 50 to 95 minutes. In particular, both high Na^+^ and high Mg^2+^ levels reduced growth by a third. Despite this substantial effect on growth, we observed no significant changes in the relative flux through different reactions in central metabolism. The only exception was a potential decrease in pentose-5-phosphate pathway use and increase in flow through malic enzyme at 300 mM Na^+^. The general picture, however, was that homeostasis in central metabolism was sufficient to ward off significant changes in relative pathway use despite large changes in overall growth rate and the pools of mRNA and proteins.

In summary, our study provides a large and comprehensive dataset for investigating the gene-regulatory response of *E. coli* under different growth conditions, both at the mRNA and the protein level. We found systematic differences in gene-expression response between exponential and stationary phase, and between mRNAs and proteins. Our dataset provides a rich resource for future modeling of *E. coli* metabolism.

## Materials and Methods

Our experimental approach was identical to the one used in our prior work on glucose starvation^10^. In particular, growth and harvesting of *E. coli* B REL606 cell pellets for the multiomic analysis was performed as previously described^10^. Similarly, after sample collection, RNA-seq, mass-spec proteomics, and metabolic flux analysis were performed as previously described^10^. For completeness, we here reproduce our complete protocol.

### Cell Growth

*E. coli* B REL606 was inoculated from a freezer stock into 10 ml of Davis Minimal medium supplemented with 2 μg/l thiamine (DM)^36^ and limiting glucose at 500 mg/l (DM500) in a 50 ml Erlenmeyer flask. This culture was incubated at 37°C with 120 r.p.m. orbital shaking over a diameter of 1". After overnight growth, 500 μl of the culture was diluted into 50 ml of prewarmed DM500 in a 250 ml flask and grown for an additional 24 h under the same conditions. On the day of the experiment, 500 μl of this preconditioned culture was added to ten 250 ml flasks, each containing 50 ml DM500, to initiate the experiment. At each time point, aliquots of these cultures were removed as necessary to harvest a constant number of cells given the changes in cell density over the growth curve. Each sample was pelleted by centrifugation, washed with sterile saline (0.85% (w/v) NaCl), and then spun down again. After removing the supernatant, the resulting cell pellet was flash frozen using liquid nitrogen and stored at −80°C. Each of the three biological replicates was performed on a separate day. Samples for each type of cell composition measurement were taken from the same batch of flasks, except for those used for flux analysis, which were grown separately in [U-^13^C] glucose. For tests of different carbon sources, the Davis Minimal (DM) medium used was supplemented with 0.5 g/L of the specified compound (glycerol, lactate, or gluconate) instead of glucose. Mg^2+^ concentrations were varied by changing the amount of MgSO_4_ added to DM media from the concentration of 0.83 mM that is normally present. For tests of different Na^+^ concentrations, NaCl was added to achieve the final concentration. The base recipe for DM already contains ~5 mM Na^+^ due to the inclusion of sodium citrate, so 95 mM NaCl was added for the 100 mM Na^+^ condition, for example. Exponential-phase samples were taken during growth when the OD_600_ reached 20-60% of the maximum achieved after saturating growth. Stationary phase samples were collected 20-24 hours after the corresponding exponential sample. The exact sampling times for each condition are provided in Supplementary Table S1.

For OD_600_ measurements, cultures were grown separately from the main batches used for harvesting cells but under identical conditions. The OD_600_ (absorbance at 600 nm) of a sample removed from the culture at each time point was measured relative to a sterile DM500 glucose blank. Doubling times were estimated from these OD_600_ measurements. Specifically, the logarithms of all OD_600_ values in the exponential part of each growth curve, defined as when OD_600_ values were between 0.05 and 0.75 times the maximum observed OD_600_ at stationary phase, were fit to a linear model with respect to time. Doubling times were calculated as log_e_2 divided by the fit slope for each biological replicate separately. Means and confidence intervals were calculated from three replicate growth curves for all conditions except for gluconate and lactate, which had measurements for only two replicates.

### RNA-seq

Total RNA was isolated from cell pellets using the RNAsnap method^37^. After extraction, RNA was ethanol precipitated and resuspended in 100 μl H_2_O. Each sample was then DNase treated and purified using the on-column method for the Zymo Clean & Concentrator-25 (Zymo Research). RNA concentrations were determined throughout the purification procedure using a Qubit 2.0 fluorometer (Life Technologies). DNase-treated total RNA (≤5 μg) was then processed with the Gram-negative bacteria RiboZero rRNA removal kit (Epicentre). After rRNA depletion, each sample was ethanol precipitated and resuspended in H_2_O again. A fraction of the RNA was then fragmented to ~250 bp using the NEBNext Magnesium RNA Fragmentation Module (New England Biolabs). After fragmentation, RNA was ethanol precipitated, resuspended in 20 μl H_2_O, and phosphorylated using T4 polynucleotide kinase (New England Biolabs). After another ethanol precipitation cleanup step, sequencing library preparation was performed using the Multiplex Compatible NEBNext Small RNA Library Prep Set for Illumina (New England Biolabs). Samples were ethanol precipitated again after library preparation and separated on a 4% agarose gel. All DNA fragments greater than 100 bp were excised from the gel and isolated using the Zymoclean Gel DNA Recovery kit (Zymo Research). Libraries were sequenced using an Illumina HiSeq 2500 at the Genomic Sequencing and Analysis Facility (GSAF) at the University of Texas at Austin to generate 2×101-base paired-end reads.

For RNA-seq analysis, we implemented a custom analysis pipeline using the REL606 *Escherichia coli* B genome (GenBank:NC_012967.1) as the reference sequence^38^. We updated annotations of sRNAs in this genome sequence using the Rfam 11.0 database^39^. Prior to mapping, we trimmed adapter sequences from Illumina reads using Flexbar 2.31^40^. Mapping was carried out in single-end mode using Bowtie2 2.2.5 with the–k 1 option to achieve one unique mapping location per read^41^. The raw number of reads mapping to each gene were counted using HTSeq 0.6.0^42^. Exact details for the full computational pipeline are available at https://github.com/marcottelab/AG3C_starvation_tc

### Proteomics

*E. coli* cell pellets were resuspended in 50 mM Tris-HCl pH 8.0, 10 mM DTT. 2,2,2- trifluoroethanol (Sigma) was added to 50% (v/v) final concentration and samples were incubated at 56°C for 45 min. Following incubation, iodoacetamide was added to a concentration of 25 mM and samples were incubated at room temperature in the dark for 30 min. Samples were diluted 10-fold with 2 mM CaCl_2_, 50 mM Tris-HCl, pH 8.0. Samples were digested with trypsin (Pierce) at 37°C for 5 h. Digestion was quenched by adding formic acid to 1% (v/v). Tryptic peptides were filtered through Amicon Ultra 30 kD spin filtration columns and bound, washed, and eluted from HyperSep C18 SpinTips (Thermo Scientific). Eluted peptides were dried by speed-vac and resuspended in Buffer C (5% acetonitrile, 0.1% formic acid) for analysis by LC-MS/MS.

For LC-MS/MS analysis, peptides were subjected to separation by C18 reverse phase chromatography on a Dionex Ultimate 3000 RSLCnano UHPLC system (Thermo Scientific). Peptides were loaded onto an Acclaim C18 PepMap RSLC column (Dionex; Thermo Scientific) and eluted using a 5-40% acetonitrile gradient over 250 min at 300 nl/min flow rate. Eluted peptides were directly injected into an Orbitrap Elite mass spectrometer (Thermo Scientific) by nano-electrospray and subject to data-dependent tandem mass spectrometry, with full precursor ion scans (MS1) collected at 60,0000 resolution. Monoisotopic precursor selection and charge-state screening were enabled, with ions of charge >+1 selected for collision-induced dissociation (CID). Up to 20 fragmentation scans (MS2) were collected per MS1. Dynamic exclusion was active with 45 s exclusion for ions selected twice within a 30 s window.

Spectra were searched against an *E. coli* strain REL606 protein sequence database and common contaminant proteins (MaxQuant using SEQUEST (Proteome Discoverer 1.4; Thermo Scientific). Fully-tryptic peptides were considered, with up to two missed cleavages. Tolerances of 10 ppm (MS1) and 0.5 Da (MS2), carbamidomethylation of cysteine as static modification, and oxidized methionine as dynamic modification were used. High-confidence peptide-spectral matches (PSMs) were filtered at <1% false discovery rate determined by Percolator (Proteome Discoverer 1.4; Thermo Scientific).

### Flux analysis

Flux ratios were obtained from the samples grown with ^13^C labeled glucose, using methods previously described^43,44^. Cell pellets were resuspended in 200 ml of 6 N HCl, hydrolyzed at 105°C overnight, and dried at 95°C for up to 24 h. To the hydrolyzed cell material we added 40 ml of dimethylformamide (DMF) and gently mixed until a “light straw” color was obtained. The DMF resuspension was transferred to a GC-MS vial with plastic insert and 40 ml of *N-tert-*butyldimethylsilyl-*N*-methyltrifluoroacetamide with 1% tert-butyldimethyl-chlorosilane (v/v); vials were capped and baked at 85°C for 2 h, and samples were analyzed within 2 days of derivitization.

Analysis of derivitized samples was performed on a Shimadzu QP2010 Plus GC-MS (Columbia, MD) with autosampler. The GC-MS protocol included: 1 mL of sample injected with 1:10 split mode at 230°C; an oven gradient of 160°C for 1 min, ramp to 310°C at 20°C/min, and hold at 310°C for 0.5 min; and flow rate was 1 mL/min in helium. A total of five runs were performed for each sample: a blank injection of DMF to waste, a blank injection of DMF to the column, and three technical replicates of each vial. Flux inference was performed using the FiatFlux software^35,44^.

### Normalization and quality control of RNA and protein counts

Our raw input data consisted of RNA and protein counts. Protein counts can be fractional, because some peptide spectra cannot be uniquely mapped to a single protein, so they are equally divided amongst these proteins. We rounded all protein counts to the nearest integer for subsequent analysis. We set the counts of all unobserved proteins to zero. For RNA, we only analyzed the counts of reads that overlapped annotated protein coding genes, i.e., reads mapping to mRNAs. This resulted in 4196 matching mRNA and protein counts for each sample. Subsequently, all mRNA and protein counts were analyzed in the same manner.

We next performed quality control, by checking replicates of the same condition for consistency. For all pairs of replicate samples, we made histograms of the log-differences of RNA or protein counts. If the two samples differ only by experimental noise, i.e., by random measurement error that causes unbiased variation in the counts of individual RNAs or proteins, then the resulting histogram should have a mode at 0 and be approximately bell-shaped. If a sample consistently shows deviations from this expectation when compared to other samples, then there are likely systematic problems with this sample. We tested the quality of our mRNA and protein samples by looking the similarity between samples collected in similar conditions but from different batches whenever possible, i.e., whenever we have at least 3 replicates. Out of 152 mRNA samples we found only two samples (samples MURI_091 and MURI_130, Supplementary Table S1) that seemed to deviate from their biological replicas. Among 105 protein samples we found no major deviation between biological replicas. Because of this broad consistency among all samples for the same growth conditions, we keep all samples for subsequent analysis.

After quality control, we normalized read counts using size-factors calculated via DESeq2^17^. Because we had many mRNAs and proteins with counts of zero at some condition, we added pseudo-counts of +1 to all counts before calculating size factors. We then used those size factors to normalize the original raw counts (i.e., without pseudo-counts).

### Clustering

We clustered normalized mRNA and protein counts based on their Euclidian distance, using the complete linkage method implemented in the `flashclust`^45^ package, which is a faster implementation of the `hclust` function in R. This method defines the cluster distance between two clusters as the maximum distance between their individual components^46^. At every stage of the clustering process, the two closest clusters are merged into the next bigger cluster. The final outcome of this process is a dendogram that measures the closeness of different samples to each other.

To assess whether the clustering process significantly grouped similar samples together, we employed a reshuffling test. For any category that we tested for significant clustering (e.g., carbon source, Na stress, or batch number), we calculated the mean cophenetic distance in the clustering dendogram between all pairs belonging to the same level of the categorical variable tested (e.g., same carbon source). We then repeatedly reshuffled the labeling within each category and recalculated the mean cophenetic distance each time. Finally, we calculated *z* scores of the original cophenetic distance relative to the distribution of reshuffled values.

### Identifying differentially expressed genes

We used DESeq2^17^ to identify differentially expressed mRNAs and proteins across conditions. We used two reference conditions in our comparisons, one for exponential phase and one for stationary phase. The reference conditions always had glucose as carbon source and base Na+ and Mg^2^+ concentrations. We did not compare exponential phase to stationary phase samples, since this comparison was done in depth previously^10^ for samples grown on glucose and with base Na+ and Mg^2^+ concentrations.

We corrected for possible batch effects by including batch number as a predictor variable in the design formula of DESeq2. In general, our design formula was `~batch_number + variable_of_interest, where variable_of_interest` was either a categorical variable representing the carbons source or growth phase (exponential or stationary) or a quantitative variable representing Na^+^ level or Mg^2+^ level. In analyses controlling for growth rate in addition to batch effects, our design formula was `~batch_number + doubling_time + variable_of_interest`.

We considered genes as differentially expressed between two conditions if their log_2_ fold change was > 1 and their FDR-corrected *P* value < 0.05. We subsequently annotated differentially expressed genes with DAVID^20^ version 6.8 Beta released in May 2016. We considered both KEGG pathways^18^ and GO annotations^19^.

### Statistical analysis and data availability

All statistical analyses were performed in R. All processed data and analysis scripts are available on github: https://github.com/umutcaglar/ecoli_multiple_growth_conditions Raw Illumina read data and processed files of read counts per gene and normalized expression levels per gene have been deposited in the NCBI GEO database^47^ (accession GSE67402 for the glucose time-course previously published^10^, accession GSE94117 for all other experiments). The mass spectrometry proteomics data have been deposited to the ProteomeXchange Consortium via the PRIDE partner repository^48^ (accession PXD002140 for the glucose time-course previously published^10^, accession PXD005721 for all other experiments). Raw GC-MS data for flux measurements have been deposited on the Texas Data Repository Dataverse at http://dx.doi.org/10.18738/T8/UG3TUR.

## Acknowledgments

This study was funded by Army Research Office (ARO,http://www.arl.army.mil/) grant W911NF-12-1-0390 to CJM, EMM, JEB, and COW. EMM also acknowledges support from the NIH (DP1 OD009572) and Welch Foundation (F1515). COW also acknowledges support from the NIH (R01 GM088344, R01 AI120560) and the NSF (Cooperative agreement no. DBI-0939454, BEACON Center). The Texas Advanced Computing Center (TACC) at The University of Texas at Austin provided high-performance computing resources.

## Contributions

M.U.C., J.R.H., C.J.M., E.M.M., J.E.B., C.O.W. conceived the study and designed the experiments. J.R.H., C.S.B., D.R.B., S.M.C., A.D. performed the experiments. V.S., D.K.S. contributed computer code used for data analysis. M.U.C., J.H.R., W.F.L., B.L.S., V.S., D.V.W., J.E.B., C.O.W. analyzed the data. M.U.C., W.F.L., V.S., D.V.W. prepared the figures. M.U.C., B.L.S., D.V.W., C.O.W. wrote the initial paper draft. All authors reviewed and edited the final manuscript

